# Multiplexed conditional genome editing with Cas12a in *Drosophila*

**DOI:** 10.1101/2020.02.26.966333

**Authors:** Fillip Port, Maja Starostecka, Michael Boutros

## Abstract

CRISPR-Cas genome engineering has revolutionised biomedical research by enabling targeted genome modification with unprecedented ease. In the popular model organism *Drosophila melanogaster* gene editing has so far relied exclusively on the prototypical CRISPR nuclease Cas9. The availability of additional CRISPR systems could expand the genomic target space, offer additional modes of regulation and enable the independent manipulation of genes in different cell populations of the same animal. Here we describe a platform for efficient Cas12a gene editing in *Drosophila*. We show that Cas12a from *Lachnospiraceae bacterium*, but not *Acidaminococcus spec*., can mediate robust gene editing *in vivo*. In combination with most crRNAs, LbCas12a activity is strongly suppressed at lower temperatures, enabling control of gene editing by simply modulating temperature. LbCas12a can directly utilize compact crRNAs arrays that are substantially easier to construct than Cas9 sgRNA arrays, facilitating multiplex genome engineering of several target sites in parallel. Targeting genes with arrays of three crRNAs results in the induction of loss-of function phenotypes with comparable efficiencies than a state-of-the-art Cas9 system. Lastly, we show that cell type-specific expression of LbCas12a is sufficient to mediate tightly controlled gene editing in a variety of tissues, allowing detailed analysis of gene function in this multicellular organism. Cas12a gene editing substantially expands the genome engineering toolbox in this organism and will be a powerful method for the functional annotation of the *Drosophila* genome. This work also lays out principles for the development of multiplexed transgenic Cas12a genome engineering systems in other genetically tractable organisms.

## Introduction

Understanding the development of multicellular organisms and the mechanisms that govern their health and disease requires detailed knowledge of the underlying genetic loci. Research in genetically tractable model organisms has been particularly powerful for functional genome annotation as it allows to experimentally introduce genetic variants and observe the resulting phenotype. Chemical, biological and physical methods to induce mutations in the genome have been used for many decades, but the fact that these methods introduce genetic variation at random makes systematic gene discovery inefficient. As a result, functionally characterized genetic alleles have so far been described only for a minority of genes (Kaufman 2017). RNAi constructs for gene knock-down have partially filled this gap, but are themselves limited by off-target effects and residual gene expression (Perkins et al. 2015). The efficient targeted introduction of mutations at specific loci would facilitate a more systematic assessment of the functional roles of genomic elements, but methods for such genome engineering have become broadly accessible only relatively recently. This has been largely driven by the discovery that the prokaryotic clustered regularly interspersed short palindromic repeat (CRISPR) - CRISPR associated (Cas) system can be easily programmed to induce DNA double strand breaks (DSBs) at defined loci in the genome of a large variety of organisms (Knott and Doudna 2018; Jinek et al. 2012; Carroll 2014).

In the widely used model organism *Drosophila melanogaster* CRISPR-Cas genome engineering has so far almost exclusively relied on the use of the RNA-guided DNA endonuclease Cas9 from *Streptococcus pyogenes* (SpCas9) (Bier et al. 2018). When expressed from a stable transgenic source SpCas9 mediates highly efficient induction of targeted DSBs in the *Drosophila* genome, which often lead to small insertions or deletions (Indels) at the target locus or can be harnessed for the precise introduction of novel sequences through homology directed repair (Port et al. 2014; 2020; Gratz et al. 2014; Kondo and Ueda 2013). However, the use of Cas9 for mutagenesis is limited to target sites adjacent to a cognate protospacer adjacent motif (PAM), which in the case of SpCas9 is NGG (Jinek et al. 2012). Furthermore, Cas9-induced mutations are predominantly small (1-10 bp) deletions, which when occurring in-frame often do not disrupt gene function. In addition, a single genome engineering system is not sufficient to independently engineer target loci in different cell types of the same organism, a prerequisite to take full advantage of an *in vivo* model to study complex organismal biology. Therefore, there exists a need for additional genome engineering systems in *Drosophila*, which can be used instead or in parallel to Cas9.

CRISPR-Cas12a (formerly known as Cpf1) is a class 2 type V CRISPR system that has been adopted for genome engineering in several organisms (Zetsche et al. 2015; Moreno-Mateos et al. 2017; Malzahn et al. 2019; Chow et al. 2019). Cas12a is a RNA-guided DNA endonuclease with a number of attributes distinct from Cas9, making it an interesting candidate to complement the CRISPR toolbox also in *Drosophila*. First, the target space for Cas12a mediated genome editing is non-overlapping with that of Cas9, as it requires a TTTV PAM (Zetsche et al. 2015; H. K. Kim et al. 2017). Second, Cas12a mediates a staggered DSB, which has been suggested to lead to distinct DNA repair outcomes (Zetsche et al. 2015; Swarts and Jinek 2019). Third, Cas12a only requires a short CRISPR RNA (crRNA) for gene targeting and has RNase activity to process single crRNAs from a crRNA array, a potential advantage for multiplex genome engineering (Fonfara et al. 2016; Zetsche et al. 2017; Campa et al. 2019). The most commonly used Cas12a enzymes originate from *Acidaminococcus* (AsCas12a) and *Lachnospiraceae bacterium* (LbCas12a) and both enzymes function with comparable efficiency in mammalian cells (Zetsche et al. 2015; Wang et al. 2018). We previously assessed Cas12a performance in *Drosophila* focused exclusively on AsCas12a and reported low activity compared to SpCas9 (Port and Bullock 2016). A subsequent study in zebrafish and *Xenopus* demonstrated that Cas12a activity is strongly temperature-dependent and showed that AsCas12a has low activity at temperatures below 30°C (Moreno-Mateos et al. 2017). This is likely to limit the utility of AsCas12a in *Drosophila*, which do not tolerate long-term exposure to temperatures above 30°C. However, in fish and frogs LbCas12a was found to have higher activity at low temperatures, raising the possibility that LbCas12a might efficiently function as RNA-guided DNA endonuclease also in flies (Moreno-Mateos et al. 2017).

Here, we report efficient multiplexed and conditional Cas12a genome engineering in *Drosophila*. We show that LbCas12a, in contrast to AsCas12a, mediates robust mutagenesis of endogenous target genes. With most, but not all, crRNAs, LbCas12a activity is strongly influenced by temperature, being inactive at 18°C and becoming active at 29°C, and thus enabling temperature controlled genome editing. We also show that LbCas12a can utilize compact crRNA arrays for multiplex genome engineering *in vivo* and using three crRNAs per target gene results in gene disruption with similar efficiency than SpCas9. Lastly, we demonstrate that tissue-specific expression of LbCas12a is sufficient for spatially restricted mutagenesis *in vivo*. The tools presented here substantially expand the genome engineering toolbox in *Drosophila* and enable targeting of previously inaccessible genomic target sites, facilitate simultaneous mutagenesis of more than two loci to pave the way for the independent manipulation of different cell populations in the same organism.

## Results and Discussion

### Temperature controlled Cas12a gene editing

In order to systematically characterize LbCas12a and AsCas12a for genome engineering applications in *Drosophila*, we established a toolbox for the expression of each Cas12a enzyme and its cognate crRNAs *in vivo* (Fig. 1A, B). We previously described a plasmid for the expression of AsCas12a crRNAs (*pCFD7*) and a transgenic fly line expressing AsCas12a under the ubiquitous *act5C* promoter (Port and Bullock 2016). We generated an equivalent expression plasmid for LbCas12a crRNAs (*pCFD8*) and an *act5c-LbCas12a* transgenic fly line, inserted at the same genomic landing site than the AsCas12a transgene (Fig. 1B). Plasmids *pCFD7* and *pCFD8* both express crRNAs from the strong, ubiquitous *U6:3* promoter, but differ in the repeat-derived 5’ stem-loop of the crRNA, which is unique for each enzyme. Importantly, AsCas12a and LbCas12a require the same PAM sequence (Zetsche et al. 2015), enabling direct comparisons of the two enzymes using crRNAs targeting the exact same target site.

**Figure 1:**
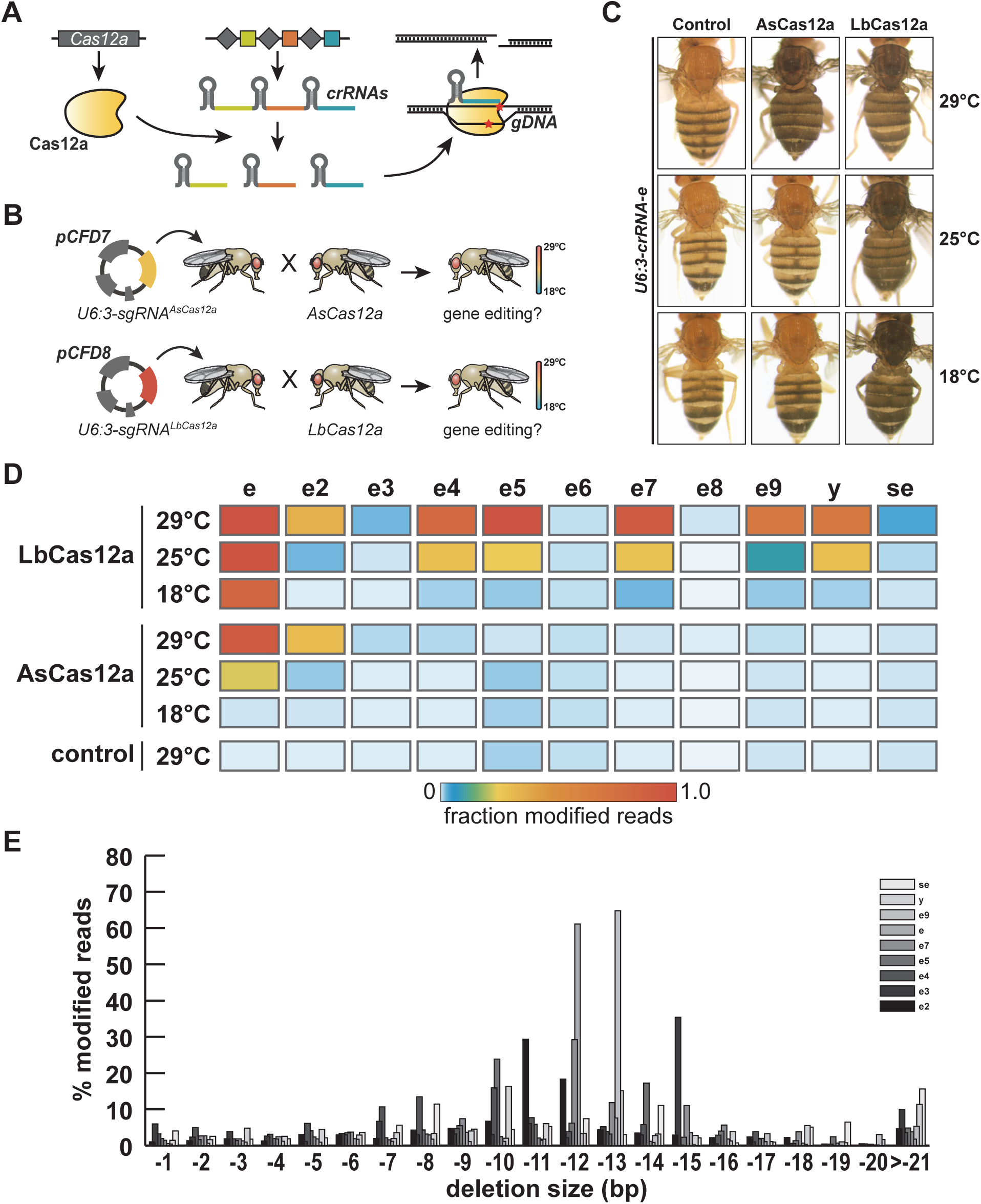
LbCas12a mediates efficient temperature-sensitive gene editing in *Drosophila*. (A) Schematic of the Cas12a genome engineering system. Cas12a pocesses RNAse activity and can process its own crRNA arrays. Upon on-target binding Cas12a cuts dsDNA, producing ends with 5’ overhangs. (B) Schematic of the experimental workflow. Cas12a crRNAs are cloned into expression vector pCFD7 (for AsCas12a) or pCFD8 (for LbCas12a). Plasmids are integrated at defined genomic locations and crossed to the respective nuclease. Offspring inheriting Cas12a and crRNA transgenes are raised at different temperatures and assessed for on-target gene editing. (C) Cas12a mediated mutagenesis of *ebony* (*e*) at different temperatures. Images of flies expressing *crRNA-e* and either AsCas12a or LbCas12a and raised at the indicated temperatures are shown. Disruption of *e* results in dark coloration of the cuticle. AsCas12a mediates efficient mutagenesis only at 29°C, while LbCas12a mediates gene disruption at all tested temperatures. (D) Assessment of gene editing efficiency by amplicon-sequencing. Activity of each nuclease was tested at 18°C, 25°C and 29°C with 11 crRNAs. Genomic target sites of each crRNA were PCR amplified and amplicon pools were subjected to deep sequencing. Gene editing activity was strongly temperature-sensitive with both nucleases. LbCas12 displayed much more robust activity, mediating strong mutagenesis with 7/11 crRNAs at 29°C, compared to 2/11 in the case of AsCas12a. (E) Size of on-target deletions mediated by LbCas12a. Shown are the distribution of deletion size in the amp-seq reads containing deletions mediated by the nine indicated crRNAs. While a large variety of deletions exist, deletions ranging between 7 and 15 bp are particularly common.

We first combined the two Cas12a enzymes with crRNAs targeting a characterized site in *ebony* (*e, crRNA-e*), which we previously found to be susceptible to AsCas12a induced gene editing in the germline with low efficiency when flies were reared at 25°C (Port and Bullock 2016). Consistent with these results *act-AsCas12a U6:3-crRNA-e* flies at 25°C had normal colouration of the cuticle, as did flies of the same genotype raised at the lower temperature of 18°C, indicating little or no inefficient disruption to *e* (Fig. 1C). In contrast, *act-AsCas12a U6:3-crRNA-e* flies reared at 29°C had dark cuticle typical of *e* loss-of function mutants, suggesting efficient mutagenesis of *e* at this higher temperature. Interestingly, flies expressing LbCas12a and crRNA-e had ebony body colouration regardless of the temperature they were raised at, suggesting that in *Drosophila* LbCas12a has higher activity at low temperatures than AsCas12a (Fig. 1C).

To evaluate if such differences in the activity of the two Cas12a enzymes at different temperatures are observed more generally, we generated 10 additional pairs of crRNA transgenes, 8 targeting additional sites in *e*, as well as one crRNA targeting *yellow* (*y*) and one crRNA targeting *sepia* (*se*) (Suppl. Table 1). We then tested these in combination with their cognate Cas12a enzyme at either 18°C, 25°C or 29°C. To quantitatively assess the amount of gene editing with high sensitivity we performed deep sequencing of PCR amplicons spanning the crRNA target sites. AsCas12a did not result in detectable gene editing at any site when flies were reared at 18°C and only mediated medium to high editing in combination with 2 of the 11 crRNAs at 29°C (Fig. 1D). In contrast, LbCas12a displayed much higher activity at 29°C, resulting in efficient gene editing when combined with 7 of the 11 crRNAs (Fig. 1D). Activity of LbCas12a was markedly reduced when animals were raised at 25°C and very low or absent at 18°C (Fig. 1D). The exception was *crRNA-e*, which mediated high mutagenesis levels at all three temperatures (Fig. 1D). Amplicon sequencing data was consistent with the level of visible phenotypes observed in adult flies (Suppl. Fig. 1). We also tested temperature dependency of Cas9-mediated gene editing with 7 sgRNAs targeting genes with visible phenotypes and did not observe any difference in phenotypic outcomes, suggesting that Cas9 functions with similar efficiency at temperatures between 18°C and 29°C (Suppl. Fig. 2). Together these experiments establish that the efficiency of Cas12a genome engineering with transgenic components in *Drosophila* is strongly temperature dependent. LbCas12a displays significantly higher activity than AsCas12a in the temperature range tolerated by *Drosophila* and in combination with many crRNAs LbCas12a can be switched from very low to high activity by increasing the temperature from 18°C to 29°C. LbCas12a is therefore a powerful CRISPR effector for inducible genome editing in *Drosophila* and we focused on this nuclease for the reminder of this study.

Targeted deep-sequencing of induced on-target edits also allowed us to profile the mutations induced by LbCas12a. Each crRNA resulted in a specific mutagenesis pattern dominated by deletions. These deletions most commonly ranged in size between 10 bp and 15bp (Fig. 1E). These results confirm and expand previous observations by us and others that Cas12a induced mutations are significantly larger than those induced by Cas9 (Port and Bullock 2016; D. Kim et al. 2016; Bernabé-Orts et al. 2019). Larger deletions are often desirable as they have a higher chance to impair the function of genetic elements.

### Multiplexed gene editing with crRNA arrays

Cas12a has been shown to possess RNAse activity and be able to excise single crRNAs from a multiplexed crRNA array (Fonfara et al. 2016). This substantially simplifies multiplex gene editing compared to the Cas9 system, which requires additional functional elements, such as tRNAs or ribozymes, to produce multiple sgRNAs from a single transcript (Xie, Minkenberg, and Yang 2015; Nissim et al. 2014). Furthermore, Cas12a crRNAs are significantly smaller than Cas9 sgRNAs, which facilitates the cloning of such multiplexed arrays. To test if LbCas12a is able to simultaneously edit several target sites in the *Drosophila* genome in combination with an array of crRNAs encoded on the same transcript, we generated transgenic flies harbouring an array of three crRNAs targeting the Wnt secretion factor *evenness interrupted* (*evi*/*wls*). Transgenic *U6:3-crRNA-evi*^*3x*^ flies were crossed to *act-LbCas12a* flies and incubated at 29°C. The offspring died at pupal stage, the same phenotype as *evi* null mutant animals (Bänziger et al. 2006; Bartscherer et al. 2006), indicating efficient gene disruption. To confirm that lethality was caused by disruption of *evi* and to test which crRNAs in the array were active, we extracted genomic DNA from *act-LbCas12a pCFD8-crRNA-evi*^*3x*^ pupae, amplified the *evi* locus and cloned and sequenced 19 individual PCR products. 17 PCR products contained mutations at one or several crRNA target sites or had deletions either between target sites or encompassing the entire target region (Fig. 2A). Importantly, we detected mutations at the target site of all three crRNAs, demonstrating that all crRNAs in the array were successfully employed by LbCas12a.

**Figure 2:**
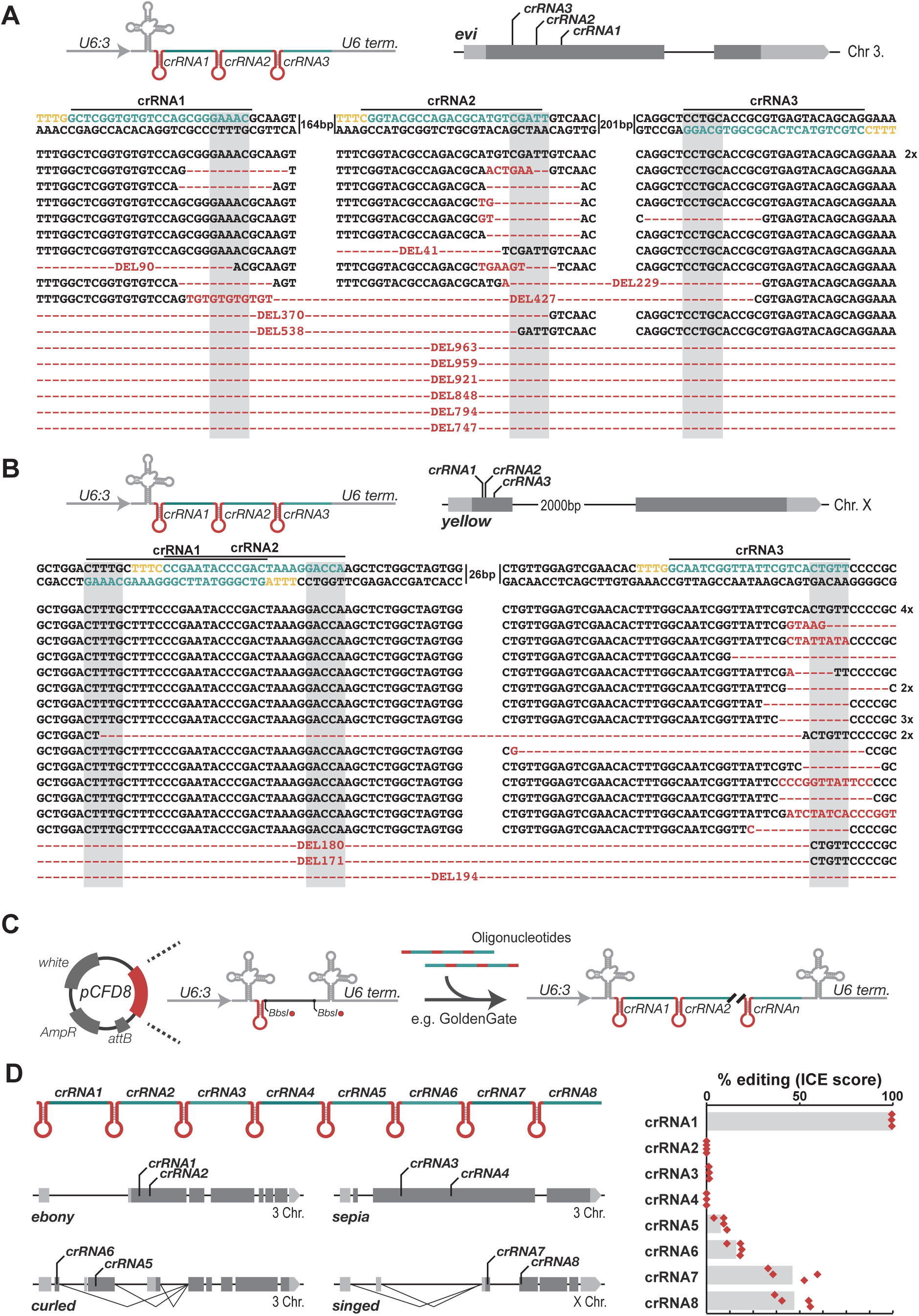
Multiplexed gene editing through crRNA arrays. (A) Gene editing of *evi* with 3 arrayed crRNAs. crRNAs were expressed from a *U6:3* promoter and combined with a *act5c-LbCas12a* transgene. Flies were raised at 29°C and died at pupal stage, indicative of efficient gene disruption. The target locus was PCR amplified from pupae and individual PCR amplicons were cloned and sequenced. 17/19 sequences were successfully edited. Edited sequences are indicated in red and the crRNA target site is indicated in green, the PAM in yellow and the window in which LbCas12a is expected to cut is shaded in gray. Mutations are detected at all target sites, demonstrating activity of all three crRNAs in the array. Indels at individual target sites are typically larger than 10bp, further supporting data presented in Fig. 1E. Deletions between individual target sites or spanning the entire locus and extending beyond the crRNA target sites (last six lines) are also detected. The size of deletions that can be detected is limited by the location of the PCR primers and deletion products might be overrepresented as small fragments are preferentially amplified in PCR and cloned more efficiently. (B) Gene editing of *y* with three arrayed crRNAs. The experiment was conducted as described above and *act-LbCas12a u6:3-crRNA-y*^*3x*^ flies had yellow cuticles indicating successful gene disruption. Sequencing of PCR amplicons revealed mutations in 21/25 amplicons, with highly efficient gene editing detected at the target site of the third crRNA in the array and editing with low efficiency at the target site of the first crRNA. No mutations were detected at the target site of the second crRNA. (C) Due to the compact size of Cas12a crRNAs several can be encoded on commercially available oligonucleotides and oligos can be fused to construct larger crRNA arrays encoded in *pCFD8*. (D) Multiplex gene targeting of four genes with two crRNAs each. Flies transgenic for the 8 x crRNA array and *act5C-LbCas12a* were raised at 29°C and had ebony cuticles and bristles displaying the singed phenotype. Each target locus was PCR amplified and amplicons were subjected to Sanger sequencing. Efficiency of gene editing was inferred from sequencing traces by ICE analysis (see methods). High levels of activity were detected for crRNA1 targeting *e*, crRNAs 7 and 8 had intermediate activity and crRNAs 5 and 6 showed low to medium activity. crRNAs 2-4 were inactive. crRNAs 1 (crRNA-e in Fig. 1C and D), 2 (crRNA-e6) and 4 (crRNA-se) were in parallel also tested when expressed as single crRNAs in *pCFD8* (see Fig. 1D) with consistent results.

We also generated a crRNA array encoding three crRNAs targeting *y*. In combination with *act5C-LbCas12a* this lead to the development of yellow cuticle in flies raised at 29°C. Sequencing of the target locus revealed mutation in 21 of 25 sequenced PCR amplicons, with the third crRNA being highly active, the first crRNAs having low activity and no mutations being detected at the target site of the second crRNA (Fig. 2B).

Next, we set out to test if a larger crRNA array would also be efficiently utilised by LbCas12a and could be used to edit several genes in parallel. Creation of larger crRNA arrays is significantly helped by the fact that three crRNAs can be encoded on commercially available oligonucleotides and oligos can be efficiently assembled into crRNA arrays in *pCFD8* (Fig. 2C). We generated a *pCFD8* plasmid containing eight crRNAs, targeting the genes *e, se, curled* (*cu*) and *singed* (*sn*) with two crRNAs per gene. We crossed *pCFD8-crRNA-e*^*2x*^*-se*^*2x*^*-cu*^*2x*^*-sn*^*2x*^ transgenic flies to *act-LbCas12a* flies and incubated the flies at 29°C. The offspring had ebony cuticle and bristles displaying the *sn* loss-of function phenotype, but did not have sepia eye coloration or curled wings. To directly monitor the editing efficiency at each target site, we extracted genomic DNA, amplified the target sites and sequenced the resulting PCR amplicons. Deconvolving the Sanger sequencing chromatograms (see methods) revealed that 3 target sites were edited with high efficiency, 2 sites with low efficiency and editing was undetectable at 3 sites (Fig. 2D). Of the three inactive crRNAs two were also tested as individual crRNAs in *pCFD8* and also showed no activity in this configuration (Fig. 1D, Suppl. Table 1), suggesting that low activity in these cases is intrinsic to the crRNA and not caused by their presence in a larger array.

Together these results demonstrate that in *Drosophila* LbCas12a can directly utilise arrays of several crRNAs to mediate multiplexed genome editing.

### Comparing the robustness of LbCas12a and Cas9

The experiments described so far establish LbCas12a as a potent genome engineering tool in *Drosophila*. However, the robustness of the Cas12a system, i.e. the fraction of crRNAs mediating efficient editing of their target site, appears lower than what is typically observed with transgenic SpCas9 systems (Fig. 1D) (Port, Muschalik, and Bullock 2015; Port et al. 2020; Kondo and Ueda 2013). We hypothesized that this could potentially be compensated for by a larger degree of crRNA multiplexing, which is facilitated by the small size of such arrays. A direct comparison of the activity of Cas9 and Cas12a at the same is not possible due to their different PAM requirements and crRNA topologies. Instead, we approached this question from a practical standpoint, comparing a Cas12a system with crRNA arrays that are easily and economically generated with a publically available, state-of-the-art resource for Cas9-mediated gene disruption.

We generated LbCas12a crRNA arrays by pooled one-step cloning, each encoding 3 crRNAs targeting the same gene at independent positions. These were stably integrated into the *Drosophila* genome at a defined position and combined with *act-LbCas12a* through a genetic cross (Fig. 3A). We then compared the phenotypic outcome of CRISPR-Cas12a mutagenesis with experiments set up in parallel utilizing the Heidelberg CRISPR Fly Design (HD_CFD) library, a large-scale collection of transgenic Cas9 sgRNA pairs (Port et al. 2020). All experiments were performed at 29°C. In total, we targeted 39 genes, each encoding a protein kinase, with either CRISPR-Cas12a or CRISPR-Cas9. Of these, 23 are described to be essential for *Drosophila* development, 2 have been shown to lead to lethality with incomplete penetrance (referred to as ‘semi-lethal’) when inactivated, 7 are established to be non-essential and for the remaining 7 only limited information is available about their loss-of function phenotype (see methods, Fig. 3B).

**Figure 3:**
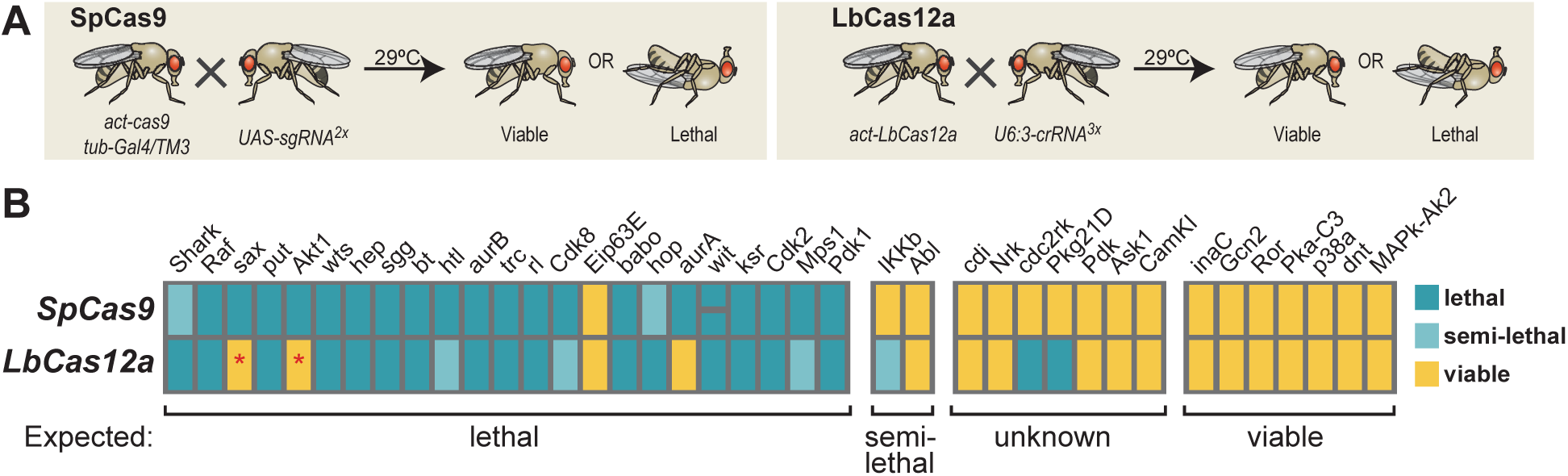
Phenotypic comparison of LbCas12a and SpCas9 gene editing. (A) Schematic of the experimental set-up. Gene targeting by SpCas9 was performed using sgRNA transgenes of the Heidelberg CRISPR Fly Design library, which encode two sgRNAs expressed from the Gal4/UAS system. LbCas12a gene editing was induced with crRNA arrays encoding three crRNAs per gene encoded in *pCFD8*. Flies ubiquitously expressing either nuclease and the respective crRNA or sgRNAs were raised at 29°C and lethality was scored 14-17 days after egg laying. (B) SpCas9 and LbCas12a gene editing both robustly identify essential and non-essential genes. Summary of the observed phenotypes is shown, with genes grouped according to the phenotype expected based on existing literature (noted below each group). Semi-lethal refers to lethality with incomplete penetrance, where between 30-70% of animals do not survive. Targeting of *sax* and *Akt1* with LbCas12a resulted in viable flies with specific morphological phenotypes, which indicated that gene targeting was successful, but not sufficiently efficient to cause lethality (red asterisk). Targeting of *wit* with SpCas9 was performed with two independent sgRNA transgenes, which both resulted in lethality.

Remarkably, CRISPR-Cas9 mutagenesis with HD_CFD sgRNAs targeting essential genes lead to lethality in 22/23 cases, highlighting the very high activity of this library (Fig. 3B). CRISPR-Cas12a mutagenesis led to lethality with 19/23 crRNA arrays targeting essential genes (Fig. 3B). The remaining 4 crRNA transgenes resulted in viable offspring. However, targeting of *sax* and *Akt1* resulted in specific phenotypes in the adult, indicating some on-target editing. Expression of both nucleases resulted in viable offspring when paired with crRNAs or sgRNAs targeting any of the 7 non-essential genes (Fig. 3B). Finally, both CRISPR systems resulted in viable offspring targeting most of the genes with limited information about their essentiality, except for *cdc2rk* and *Pkg21D*, which resulted in lethal offspring only when targeted by Cas12a (Fig. 3B).

Together these experiments highlight the very high efficiency and robustness of gene editing with both CRISPR systems in *Drosophila* at 29°C. While SpCas9 led to the expected phenotype when targeting 29 of the 32 genes based on previous findings, LbCas12a correctly identified 27 of the known essential or non-essential genes. Of note, gene essentiality is known to be often sensitive to genetic background effects and hence recapitulation of previously described phenotypes might not always be observed even if gene disruption was successful (Rancati et al. 2018).

### Tissue-specific Cas12a gene editing

Dissection of the often multifaceted functions of genes in multicellular organisms requires the ability to manipulate their sequence with spatial and temporal precision *in vivo*. We therefore explored whether LbCas12a could be utilised for tissue-specific mutagenesis in *Drosophila*. Conditional gene editing is typically achieved by linking the expression of CRISPR components to cis-regulatory elements with tissue-specific expression patterns. In *Drosophila* this is most commonly done via the binary Gal4-UAS system (Brand and Perrimon 1993). We generated *UAS-LbCas12a* flies and combined them with a range of Gal4 drivers, resulting in tissue-specific LbCas12a expression (Suppl. Fig. 3). We then crossed them with animals transgenic for U6:3-crRNAs to induce gene editing in Gal4 expressing cells of the offspring. First, we generated *pnr-Gal4 UAS-LbCas12a U6:3-crRNA-e* flies, which displayed dark cuticle exclusively in the *pnr-Gal4* expression domain along the dorsal midline, indicative of disruption of *e* exclusively in Gal4 expressing cells (Fig. 4A).

**Figure 4:**
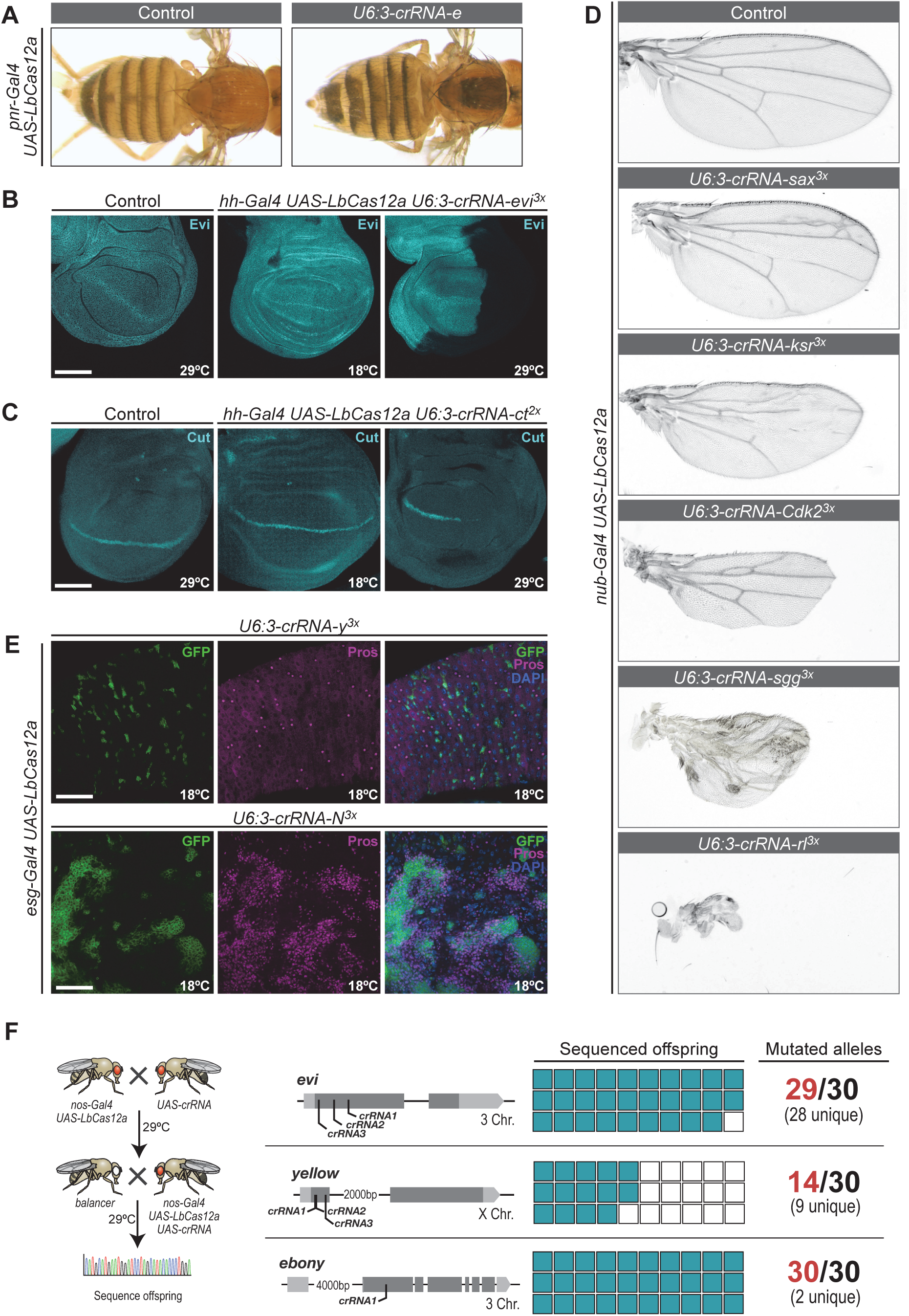
Tissue-specific gene editing with UAS-LbCas12a. (A) LbCas12a-mediated mutagenesis of *e* in the *pnr-Gal4* expression domain along the dorsal midline leads to the development of dark pigment in the cuticle. Flies were raised at 29°C. (B) Mutagenesis of *evi* in the posterior compartment of the wing imaginal disc with *hh-Gal4*. Evi protein is detected throughout the wing imaginal disc in control and *hh-Gal4 UAS-LbCas12a U6:3-crRNA-evi*^*3x*^ animals raised at 18°C, but absent from the posterior compartment in animals raised at 29°C, demonstrating efficient, temperature-controlled and spatially defined gene disruption. (C) Mutagenesis of *ct* in the posterior compartment with *hh-Gal4*. Ct protein is detected in a stripe of cells in control and *hh-Gal4 UAS-LbCas12a U6:3-crRNA-evi*^*3x*^ animals raised at 18°C, but absent from the posterior compartment in animals raised at 29°C, demonstrating efficient, temperature-controlled and spatially defined gene disruption. (D) Tissue-specific mutagenesis of essential genes in the wing primordium with *nub-Gal4*. Wings from animals expressing *nub-Gal4 UAS-LbCas12a* and the indicated crRNA array are shown. When combined with ubiquitous *act5C-LbCas12a* these crRNAs lead to lethality, but tissue-restricted expression with *nub-Gal4* gives rise to viable or semi-viable flies. Wings of such animals display specific and informative phenotypes affecting wing size, vein formation, bristle formation or wing margin formation. (E) Intestinal tumorigenesis induced by conditional LbCas12a mutagenesis of *N. esg-Gal4* was used to express LbCas12a in adult intestinal stem cells. *esg-Gal4 UAS-LbCas12a U6:3-crRNA-N*^*3x*^ animals raised at 29°C are lethal before adulthood, consistent with the known expression of *esg-Gal4* in larval and pupal tissues. Reducing the culture temperature to 18°C leads to viable adults that develop intestinal tumors characterized by expression of *pros*, a hallmark of tumors caused by loss of *N*. Mutagenesis of *y* was used as a control, as *y* does not play a role in intestinal stem cells. (F) Efficient Cas12a-mediated mutagenesis in the germline to create heritable alleles.

Next, we attempted conditional mutagenesis in wing imaginal discs. We used *hh-Gal4 UAS-LbCas12a* to knock-out *evi* selectively in the posterior compartment and analysed the outcome by directly visualising Evi protein. Evi is ubiquitously expressed in wildtype tissue, but was undetectable in the P-compartment of wing discs of *hh-Gal4 UAS-LbCas12a U6:3-crRNA-evi*^*3x*^ animals raised at 29°C (20/22 discs, Fig. 4B). Wing discs of animals of the same genotype raised at 18°C expressed Evi in almost all cells (Fig. 4B). In 7/13 discs we observed a weak reduction of Evi in the P-compartment, which might reflect some binding of the Cas12a-crRNA complex to the target gene (Malzahn et al. 2019). These results were further corroborated by targeting a second gene, *cut* (*ct*), a transcription factor expressed in a stripe of cells along the dorsal-ventral boundary (Fig. 4C). Mutagenesis of *ct* with *hh-Gal4 UAS-LbCas12a* at 29°C lead to lethality at the third instar larval stage. We made use of the temperature sensitivity of LbCas12a and raised animals for the first 24h at 18°C and then shifted them to 29°C. These animals survived until pupal stages, allowing us to assess *ct* expression in late third instar discs and highlighting the advantages of a temperature-switchable gene editing system. In 10 of the 13 analysed wing discs *ct* expression was completely lost exclusively in the *hh-Gal4* expression domain, while in the remaining 3 discs a few *ct* expressing cells remained (Fig. 4C). Mutagenesis was strongly suppressed in wing discs from animals raised at 18°C, where *ct* expression was unaffected in 5/9 discs and only few cells had lost *ct* in the remaining 4 samples (Fig. 4C). Together these experiments establish UAS-LbCas12a as a powerful tool for tissue-specific gene disruption. Interestingly, in all cases mutagenesis appeared strictly restricted to the *hh-Gal4* expression domain. This is in contrast to Cas9, where leaky expression of Cas9 from UAS vectors results in ectopic gene editing in combination with U6:3 driven sgRNAs (Port and Bullock 2016). Of note, the morphology of wing imaginal discs expressing an active Cas12a system was sometimes abnormal, with tissue folds forming at abnormal positions (Suppl. Fig. 4A). This was not caused by excessive amounts of cell death in these tissues (Suppl. Fig. 4B) and was independent of the target of the co-expressed crRNA. In the future, careful titration of Cas12a protein levels, as we have recently demonstrated with Cas9 (Port et al. 2020), might alleviate these effects.

We also used an additional line, *nub-Gal4 UAS-LbCas12a*, to restrict mutagenesis to the wing disc and a few other tissues and tested it with crRNA arrays that result in lethality in combination with *act5c-LbCas12a* (Fig. 3B). Adult flies emerged from crosses with 21 of the 23 tested crRNA transgenes, supporting tight restriction of gene editing to Gal4 expressing cells. Wings from such animals typically displayed highly specific phenotypes affecting their size, veins, bristles, texture or margin, which can give valuable clues about the function of the target gene (Fig. 4D).

Next, we analysed Cas12a-mediated mutagenesis in an adult tissue, the *Drosophila* midgut. Many Gal4 drivers are active at multiple stages during development, making transgene mediated gene editing of essential genes in adults challenging. We reasoned that the temperature-sensitivity of the Cas12a system could be used to modulate gene editing during different stages of development. We generated an *esg-Gal4 UAS-LbCas12a* line, which is expressed in adult intestinal stem cells, as well as various embryonic, larval and pupal tissues, and crossed it to a crRNA array targeting *Notch* (*N*). When *esg-Gal4 UAS-LbCas12a U6:3-crRNA-N* animals were reared at 29°C they died at larval to pupal stage, consistent with efficient mutagenesis of *N* in *esg* expressing cells during early development. In contrast, when animals of the same genotype were raised at 18°C they developed to adulthood, demonstrating effective suppression of mutagenesis at this temperature. However, 14 day old adults frequently displayed large tumors in their midgut, suggesting some disruption of *N* occured at this temperature (Fig. 4E). These tumors were formed by *prospero* expressing cells with a small nucleus, the known phenotype of tumors caused by loss of *N* (Ohlstein and Spradling 2006), and were not observed in control tissue (Fig. 4E). This demonstrates how temperature can be used to tune Cas12a gene editing rates to allow analysis of loss-of function phenotypes in essential genes in later stages of life.

Lastly, we explored Cas12a mediated mutagenesis restricted to the germline, an important application allowing to create heritable alleles. We generated a *nos-Gal4 UAS-LbCas12a* fly line and tested it with crRNA arrays targeting *e, y* and *evi* at 29°C. Flies expressing crRNAs targeting *evi* were viable and animals with crRNAs targeting *e* or *y* had normal colouration of their cuticle, indicative of very low or no Cas12a activity in somatic tissues (Suppl. Fig. 5). In contrast, all three crRNA arrys mediated high levels of on-target mutagenesis in the germline, as revealed by sequencing of their target sites in individual offspring (Fig. 4F). Together this demonstrates that germline-specific expression of LbCas12a allows to create heritable alleles in the germline without affecting viability of the animal.

## Conclusion

Here we present a system to perform efficient Cas12a genome engineering in a variety of tissues in *Drosophila melanogaster*. It consists of transgenic fly lines expressing LbCas12a from the ubiquitous *act5C* promoter or under the control of the conditional Gal4/UAS system and plasmids for the expression of one or several crRNAs. With the majority of active crRNAs LbCas12a is highly temperature sensitive, offering the ability to control mutagenesis through temporal regulation of the temperature at which experiments are performed. Furthermore, Cas12a targets sites orthogonal to Cas9 and results in larger mutations, which are more likely to disrupt gene function. While LbCas12a appears to function with a lower fraction of transgenic guides than SpCas9, this can be effectively compensated by a higher degree of crRNA multiplexing. As the costs of long oligonucleotides are falling and the length of oligos that are commercially available increases, it is likely that in the future larger crRNA arrays can be easily constructed, which would be expected to further increase the versatility of the Cas12a system.

We envision that in the future Cas12a will be the primary genome engineering system for AT rich loci, for applications that benefit from the possibility to control genome editing by temperature or in cases that require highly multiplexed mutagenesis. In addition, Cas12a will be used to independently confirm novel findings made with Cas9 or other technologies and in combination with Cas9 to manipulate genes in different cells of the same animal.

## Material and Methods

### Plasmid construction

Unless indicated otherwise PCRs were performed with the Q5 Hot-start 2x master mix (New England Biolabs (NEB)) and cloning was performed using the In-Fusion HD cloning kit (Takara Bio) or restriction/ligation dependent cloning. Newly introduced sequences were verified by Sanger sequencing. Oligonucleotide and gBlock sequences are listed in Supplementary Table 2.

#### act5c-LbCas12a

Plasmid act5c-LbCas12a was constructed by replacing the Cas9 open reading frame in *act5c-Cas9* (Port et al. 2014) by a *LbCas12a* open reading frame. DNA encoding LbCas12a (based on (Zetsche et al. 2015)) was ordered as gBlocks from Integrated DNA Technologies (IDT). Individual gBlocks were fused by extension PCR and PCR amplicons were purified using the Qiagen PCR purification kit according to the instructions supplied by the manufacturer. Plasmid *act5c-Cas9* was digested with EcoRI-HF and XhoI (NEB) and the plasmid backbone was gel purified. The plasmid backbone and the dsDNA fragment encoding LbCas12a were assembled by In-Fusion cloning. The proportion of the plasmid encoding LbCas12a was sequence verified by Sanger sequencing.

#### UAS-LbCas12a

Plasmid *UAS-LbCas12a* was generated by cloning *LbCas12a* encoded in *act5c-LbCas12a* into *pUASTattB* (Bischof et al. 2007). Plasmid pUASTattB was digested with EcoRI-HF and XbaI and gel purified. *LbCas12a* was PCR amplified from *act5c-LbCas12a* with primers *UASLbCas12afwd* and *UASLbCas12arev*. The purified PCR amplicon was assembled with the digested plasmid backbone using In-Fusion cloning.

#### pCFD8

Plasmid *pCFD8* was constructed as the default cloning plasmid for one or several LbCas12a crRNAs. It was generated by replacing the Cas9 sgRNA cassette in *pCFD5_w* (Addgene 112645) with the LbCas12 crRNA stem loop, a 2xBbsI restriction cassette, and a downstream tRNA. The necessity of the tRNAs in *pCFD8* has not been assessed, but previous work indicated that crRNAs from tRNA vectors have slightly higher efficiency (Port and Bullock 2016). Plasmid *pCFD5_w* was digested with BbsI-HF and XbaI and the backbone was gel purified. The LbCas12a crRNA cassette was ordered as a gBlock from IDT and assembled with the plasmid backbone using In-Fusion cloning.

#### Cloning of crRNAs into pCFD8

The compact size of Cas12a crRNA arrays allows to directly encode one or several crRNAs on commercially available oligonucleotides. A step-by-step cloning protocol will be made available at http://crisprflydesign.org. Briefly, *pCFD8* was digested with BbsI-HF and gel purified. Sequences to be introduced into *pCFD8* were ordered as complementary oligos (one for the top and one for the bottom strand) and resuspended to 100µM in ddH2O. Oligo pairs were mixed at an equimolar ratio in PCR tubes containing T4 Ligation buffer and T4 polynucleotide kinase (NEB) for 5’ phosphorylation of oligos, incubated at 37°C for 30min and then heated to 95°C and let cool down to 25°C at a rate of -3°C/min. Annealed oligos were diluted 1:200 in ddH_2_O. The oligos were designed such that after annealing the dsDNA had single-stranded overhangs complementary for the BbsI sites in *pCFD8*. The annealed oligos were ligated into the digested *pCFD8* backbone using T7 DNA ligase. Successful introduction of crRNAs was confirmed by Sanger sequencing.

### *Drosophila* strains and culture

Transgenic *Drosophila* strains used or generated in this study are listed in Supplementary Table 3. Flies were kept in incubators with 50±10% humidity with a 12h light/12h dark cycle. Temperature during the individual experiments are indicated in the figures and were regularly checked with independent thermometers.

### Transgenesis

Transgenesis was performed with the PhiC31/attP/attB system and plasmids were inserted at landing site (P{y[+t7.7]CaryP}attP40) on the second chromosome. Microinjection of plasmids into *Drosophila* embryos was carried out using standard procedures. Transgenesis of crRNA plasmids was typically performed by a pooled injection protocol, as previously described (Bischof et al. 2013). Briefly, individual plasmids were pooled at equimolar ratio and DNA concentration was adjusted to 250 ng/μl in dH_2_O. Plasmid pools were microinjected into y[1] M{vas-int.Dm}ZH-2A w[*]; (P{y[+t7.7]CaryP}attP40) embryos, raised to adulthood and individual flies crossed to P{ry[+t7.2]=hsFLP}1, y[1] w[1118]; Sp/CyO-GFP. Transgenic offspring was identified by orange eye color and individual flies crossed to P{ry[+t7.2]=hsFLP}1, y[1] w[1118]; Sp/CyO-GFP balancer flies and a stable transgenic stock was generated.

### Genotyping of crRNA flies

Individual transgenic flies from pooled plasmid injections were genotyped to determine which plasmid was stably integrated into their genome. If transgenic flies were male or virgin female, animals were removed from the vials once offspring was apparent and prepared for genotyping. In the case of mated transgenic females genotyping was performed in the next generation after selecting and crossing a single male offspring, to prevent genotyping females fertilised by a male transgenic for a different construct. The crRNA transgene was amplified by PCR from genomic DNA with primers U63seqfwd2 and pCFD8genorev2 and PCR amplicons were analysed by Sanger sequencing using primer U63seqfwd2.

### Preparation of genomic DNA

Single flies were collected in PCR tubes containing 50 µl squishing buffer (10 mM Tris-HCL pH8, 1 mM EDTA, 25 mM NaCl, 200 µg/ml Proteinase K). Flies were disrupted in a Bead Ruptor (Biovendis) for 20 sec at 30 Hz. Samples were then incubated for 30 min at 37°C, followed by heat inactivation for 3 min at 95°C. Typically 3 µl of supernatant were used in 30 µl PCR reactions.

### Quantification of gene editing efficiency

#### Estimation of editing efficiency by Sanger sequencing bulk PCR amplicons

To estimate the efficiency of gene editing by decomposition of Sanger sequencing traces, the target locus was amplified from genomic DNA by PCR. Primers (Suppl. Table 2) were typically designed to anneal approximately 300bp 5’ or 3’ of the crRNA target site. PCR amplicons were purified with the Qiagen PCR purification kit according to the instructions supplied by the manufacturer and subjected to Sanger sequencing. Sequencing chromatograms were then analysed by ICE (Inference of CRISPR Edits) analysis (Hsiau et al. 2019).

#### Estimation of editing efficiency by Sanger sequencing single PCR amplicons

To test for the presence of CRISPR edits in individual PCR amplicons the target locus was amplified as described above. After the final elongation 1 unit of Taq polymerase was added and the reaction was incubated for 10 min at 72°C to add 3’ A-overhangs. PCR amplicons were then cloned into the pCR4-TOPO vector using the TOPO TA cloning Kit for Sequencing (Life Technologies) according to the instructions by the manufacturer. DNA extracted from individual colonies was subjected to Sanger sequencing using the M13fwd primer and sequencing traces were inspected in Snapgene software.

#### Quantification of editing efficiency by amplicon-seq

For accurate and sensitive detection of on-target mutations PCR amplicons spanning the crRNA target site were sequenced on the Illumina MiSeq platform. Transgenic flies expressing the respective Cas12a enzyme under the ubiquitous *act5C* promoter were crossed to the indicated crRNA transgene and crosses were incubated at the indicated temperature (either 18°C, 25°C or 29°C). As a control *act-Cas12a* flies were crossed to transgenic flies expressing a Cas9 sgRNA, which was present in the same genetic background as the Cas12a crRNAs. Several flies of the offspring from each cross were pooled and 700 ng of genomic DNA was extracted and used as template to amplify genomic regions spanning each crRNA target site (see Suppl. Tab. 2 for primer sequences). PCR amplicons were purified with the Qiagen PCR purification kit according to the instructions supplied by the manufacturer. Amplicons were then PCR amplified using primers containing Illumina adaptors and indices. Following purification as described above, DNA concentration of each sample was measured with the Qubit 1.0 fluorometer (Thermo Fisher Scientific). Equimolar (10nM) dilutions of amplicons were pooled, 30% PhiX was added and the library was sequenced on a Illumina MiSeq (300 bp single-end reads) by the High-Throughput Sequencing Unit of the Genomics and Proteomics Core Facility (DKFZ). Obtained sequences were analyzed with CRISPResso (Pinello et al. 2016).

### Categorization of essential and non-essential genes

Target genes (Fig. 3) were categorized as essential or non-essential based on information available in Flybase (release FB2019_5). For each gene we manually curated the information available in the phenotype category. We did not consider information based on RNAi experiments, as these were typically performed with tissue-specific Gal4 drivers, and residual gene expression, which is the norm with this method, is expected to rescue lethality in some cases.

### Immunohistochemistry

Immunohistochemistry of *Drosophila* tissue was performed using standard procedures. Briefly, larva or adult intestine were dissected in ice cold PBS and fixed in either 80% ice-cold methanol in PBS for 1h on ice (for staining with anti-LbCas12a antibody) or in 4% Paraformaldehyde in phosphate buffered saline (PBS) containing 0.05% Triton-X100 for 25 min at room temperature (for staining with any of the other antibodies). Larva were washed three times in PBS containing 0.3% Triton-X100 (PBT) and then blocked for 1h at room temperature in PBT containing 1% heat-inactivated normal goat serum. Subsequently, larva were incubated with first antibody (mouse anti-LbCas12a (Sigma Aldrich) 1:20; mouse anti-Cut (DSHB, Gary Rubin) 1:30; rabbit anti-Evi (Port et al. 2008) 1:800; mouse anti-Prospero (MR1A, DSHB,C.Q. Doe) 1:1000) in PBT overnight at 4°C. The next day, samples were washed three times in PBT for 15 min and incubated for 2 h at room temperature with secondary antibody (antibodies coupled to Alexa fluorophores, Invitrogen) diluted 1:600 in PBT containing Hoechst dye. Samples were washed three times 15 min in PBT and mounted in Vectashield (Vectorlabs). To visualize apoptotic cells wing discs expressing the apoptosis sensor GC3Ai (Schott et al. 2017) were fixed in 4% PFA, washed in PBT containing Hoechst and mounted in Vectashield.

### Image acquisition, processing and analysis

Microscopy images were acquired with a Zeiss LSM800, Leica SP5 or SP8 confocal microscope in the sequential scanning mode. Image processing and analysis was performed with FIJI (Schindelin et al. 2012). Experiments were performed at least twice and more than 3 samples were analyzed for each experiment. Imaging of adult flies or wings was performed with a stereomicroscope equipped with a 14.2 Color Mosaic camera (Visitron Systems). If wings were analyzed, the flies were incubated for at least 4 hours in 50% ethanol/50% glycerol solution and individual wings were then mounted on microscopy slides in the same solution. Lightning was kept constant during image acquisition. Contrast and brightness were modified uniformly within an experiment with Photoshop or Fiji.

## Acknowledgements

We would like to thank Laura Lange and Nora Langner for excellent technical assistance and Luisa Henkel and Siamak Redhai for helpful comments on the manuscript. We are grateful for the support of the High-Throughput Sequencing Unit of the Genomics and Proteomics Core Facility and Light Microscopy Facility at DKFZ. This work has in part been supported by grants from the European Research Council (ERC) (DECODE) and the German Research Foundation (SFB1324, Z03).

## Author contributions

F.P. conceived the study, designed and performed experiments, analysed data, supervised the study and wrote the paper. M.S. designed and performed experiments, analysed data and edited the manuscript. M.B. acquired funding, supervised the study and edited the manuscript.

## Material availability

All material reported in this study is available upon request. Plasmids *pCFD8, act5c-LbCas12a* and *UAS-LbCas12a* will be available from Addgene (Addgene plasmids 140619, 140620, 140621).

