## SupplementaryMaterial for "Multiplexed conditional genome editing with Cas12a in *Drosophila*"

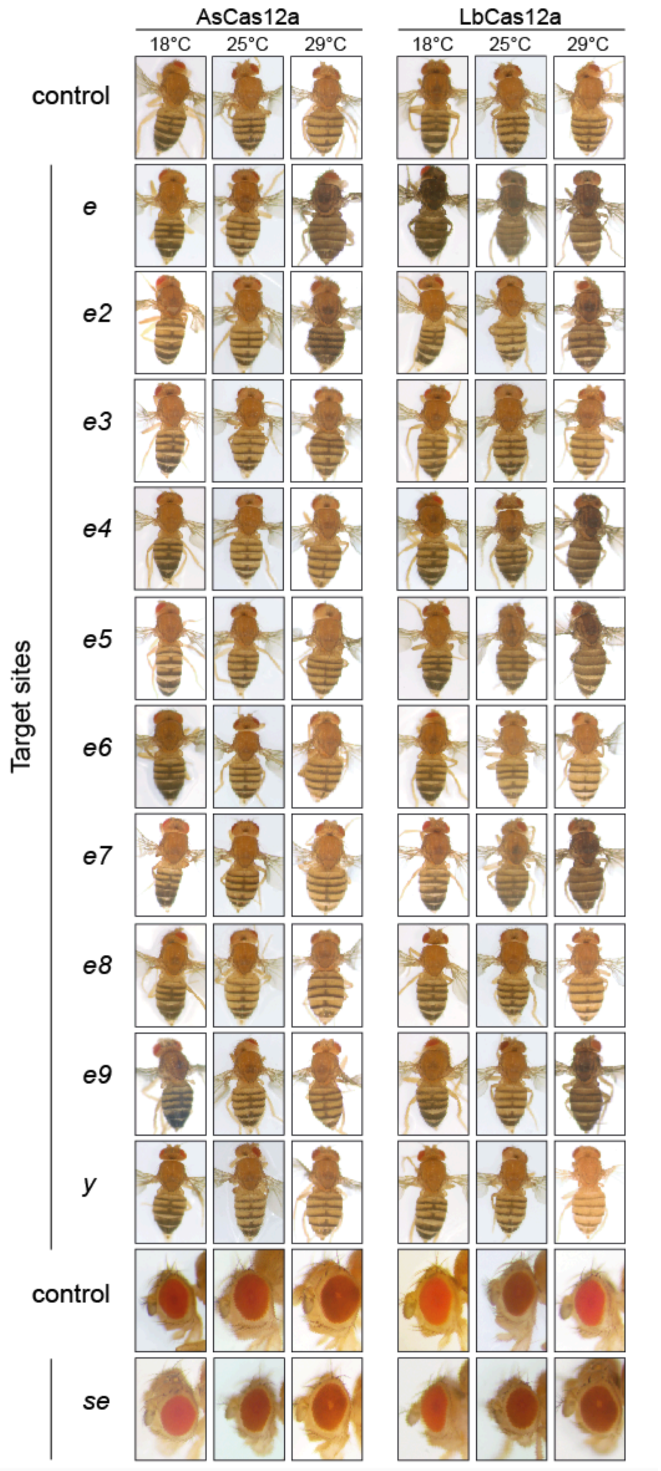


**Supplementary Figure 1: Visible phenotypes caused by AsCas12a- or LbCas12a-mediated gene editing at different temperatures.** Images of flies transgenic for the respective act5c-Cas12a and crRNA constructs incubated at the indicated temperatures. Loss-of function mutations in *e* lead to a darkening of the cuticle, while mutations in *y* result in yellow cuticle and mutations in *se* cause dark eye pigmentation (no observed). Phenotypes are observed in the same genotypes and under the same conditions that have a high proportion of mutant PCR amplicons in the amp-seq experiment shown in Fig. 1D.

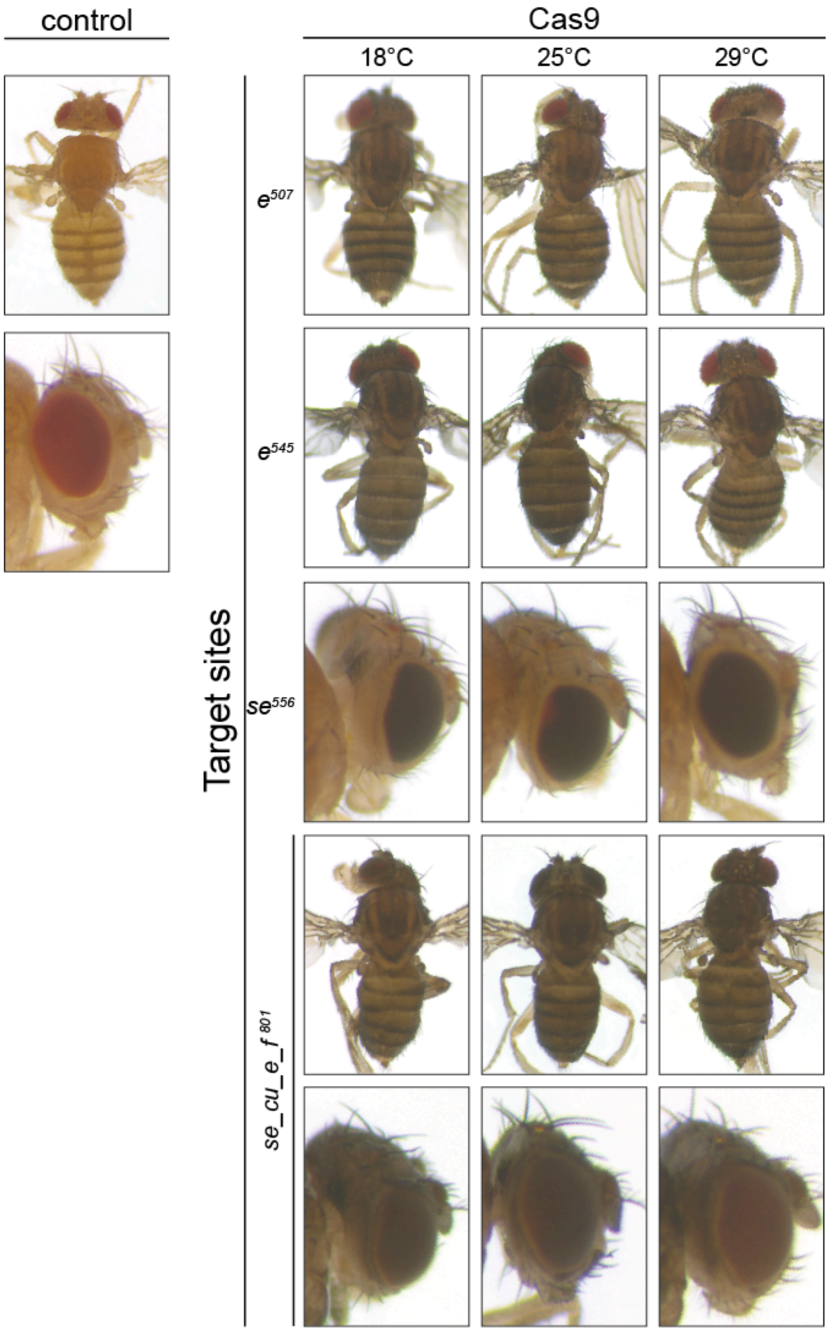


**Supplementary Figure 2: Phenotypic outcome of Cas9 gene editing at different temperatures.** Images of adult act5C-Cas9 U6:3-sgRNA flies incubated at different temperatures. Tested sgRNA transgenes were previously found to efficiently disrupt the target gene at 25℃ (Port, Muschalik, and Bullock 2015; Port and Bullock 2016). Fully penetrant phenotypes were observed at all three tested temperatures, suggesting that Cas9 has robust activity in this temperature range.

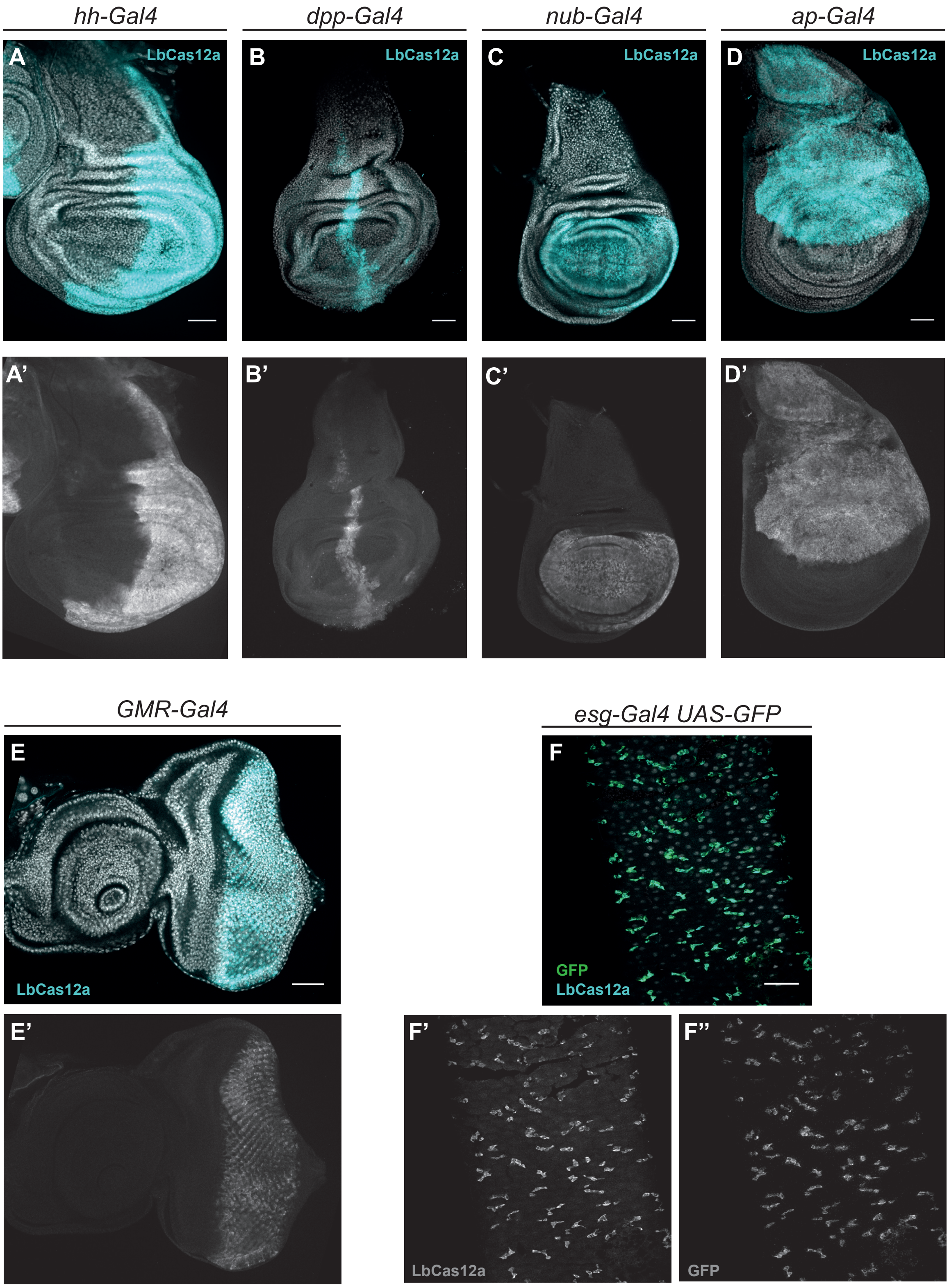


**Supplementary Figure 3: Cas12a expression patterns in various Gal4 UAS-LbCas12a lines.** Anti-LbCas12a stainings in wing imaginal discs (A - D), eye-antennal disc (E) and the adult midgut (F). LbCas12a protein is detected in the posterior compartment (*hh-Gal4*), along the anterior-posterior boundary (*dpp-Gal4*), in the wing pouch (*nub-Gal4*), the dorsal compartment (*ap-Gal4*), posterior to the morphogenetic furrow (*GMR-Gal4*) and in the intestinal stem cells (*esg-Gal4*).

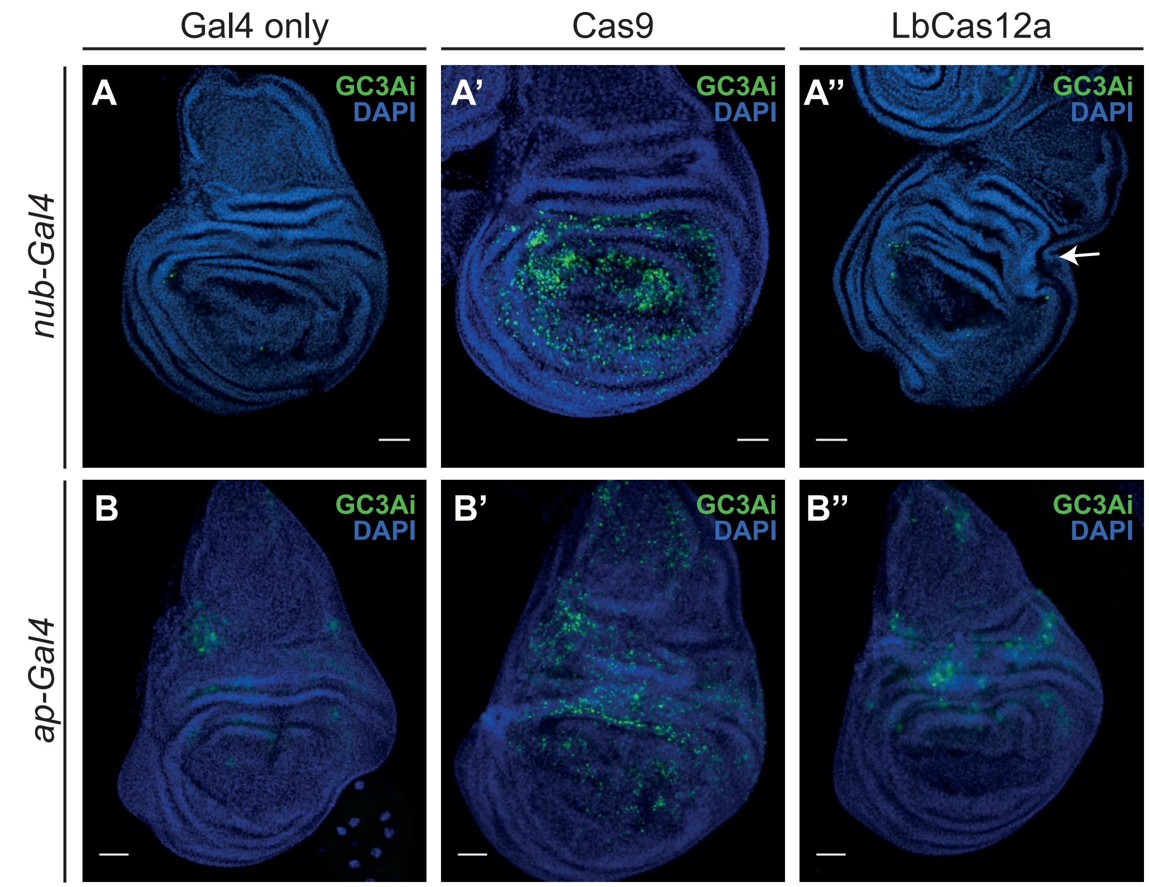


**Supplementary Figure 4: Cas12a overexpression sometimes causes morphological abnormalities, but no clear increase in the number of apoptotic cells.** Wing imaginal discs expressing the indicated Gal4 driver and Cas nuclease and the apoptosis sensor GCA3i (Schott et al. 2017). Wing discs expressing LbCas12a are often smaller than age matched controls and sometimes show excessive tissue folding (arrow). However, while expressing high levels of Cas9 nuclease results in a clear increase in the number of apoptotic cells in the wing disc, no such increase was observed when LbCas12a was expressed.

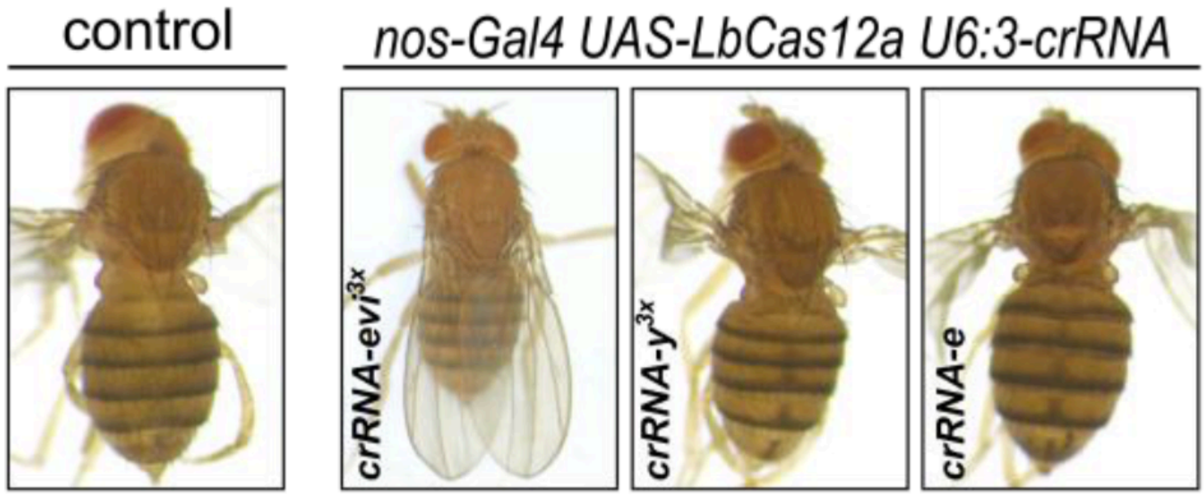


**Supplementary Figure 5: Flies expressing LbCas12a under *nos-Gal4* show no signs of mutagenesis in somatic cells.** Images of adult flies expressing *nos-Gal4 UAS-LbCas12a*

and the indicated U6:3-crRNA transgene are shown. Crosses were performed at 29°C and resulted in efficient mutagenesis of the target genes in the germline (Fig. 4F). Flies harboring a U6:3-crRNA$^{evi}$ are viable and morphologically normal, indicating little or no somatic mutagenesis of *evi*. Furthermore, flies harboring crRNA transgenes targeting *e* or *y* have normal coloration of the cuticle, indicating that mutagenesis is tightly restricted to the germline.

**Supplied as separate files:**

**Supplementary Table 1:** Target sites of individual crRNAs used in this study.

**Supplementary Table 2:** Sequences of oligonucleotide used in this study.

**Supplementary Table 3**: Fly lines used in this study.
